# Adaptive Prediction: The Brain Trades Phonemic for Semantic Expectations Under Acoustic Uncertainty

**DOI:** 10.1101/2025.07.18.665546

**Authors:** Giorgio Piazza, Marco Sala, Rebecca Guerrini, Martin M. Winchester, Francesca Peressotti

## Abstract

Listeners daily adapt to new talkers, yet how acoustic variability challenges the brain’s predictive mechanisms during speech processing remains unclear. Here, we used EEG and Temporal Response Function to examine neural responses to continuous speech narrated by a single talker (Single), enabling stable acoustic model formation, or multiple talkers (Multi), introducing acoustic uncertainty. We assessed whether talker variability influences phoneme recognition and predictive processing, indexed by neural responses to phonemes, phonemic and semantic surprisal. In the Multi condition, responses to phonemes increased but responses to phonemic surprisal decreased, indicating greater speech perception demands and weaker phonemic predictions. Conversely, semantic surprisal responses were stronger, suggesting increased reliance on lexical-semantic predictions. These findings reveal a trade-off in the brain’s predictive mechanisms, where acoustic uncertainty reduces lower-level phonemic anticipation but promotes higher-level semantic prediction. Adaptive processing underscored the brain’s ability to dynamically adjust predictions across linguistic levels, promoting speech comprehension in variable environments.

**Significance:** Humans attend to many different talkers daily, switching between them apparently without any effort. This stems from the brain’s ability to construct adaptive, probabilistic models of speech. In this study, we provide novel evidence that predictions in language comprehension are sensitive to acoustic uncertainty. Specifically, under conditions of increased acoustic uncertainty, listeners rely more on bottom-up information in phoneme recognition and reduce the anticipation of phonemic information. This is accompanied by enhanced semantic prediction, suggesting that listeners compensate for acoustic uncertainty by increasing reliance on higher-level contextual representations. This flexible approach allows for robust comprehension, highlighting the brain’s capacity to dynamically adjust its predictive processing to accommodate varying acoustic environments.

**Teaser:** Talker-driven acoustic uncertainty reduces phoneme predictions but boosts semantic inference.

## 1. Introduction

Humans interact daily with different talkers, many of whom are unfamiliar and vary widely in how they produce speech. For example, during a conference question time, listeners may be exposed to a rapid succession of unfamiliar talkers that differ in accent, voice quality, fundamental frequency, speech rate, and articulation style. Such variability poses a fundamental challenge of spoken language comprehension: the acoustic signal is continuous and highly variable, and there is no one-to-one mapping between its physical properties and the abstract linguistic categories it conveys, such as phonemes or words. Yet, listeners typically comprehend speech with remarkable ease. One influential account holds that comprehension is not purely reactive but predictive, with listeners continuously generating probabilistic expectations about upcoming input (1, 2). These predictions are thought to operate across multiple levels of representation, from phonemes to words and sentences. However, most evidence for predictive processing has been obtained under conditions that minimize variability in the acoustic signal, typically using a single talker and/or highly controlled speech (3, 4). As a result, it remains unclear whether and to what extent predictive mechanisms are robust to the fine-grained acoustic variability that characterises natural listening.

This gap is critical because prediction is assumed to operate probabilistically: the brain continuously forms graded expectations reflecting the inherent uncertainty in both the speech signal and the linguistic content (2, 5, 6). In speech perception, phonemes are not fixed entities but distributions across multiple acoustic dimensions that vary across talkers (5, 7). Listeners can adapt to this variability by constructing talker-specific distributional models of the acoustic signal, enabling predictions based not only on what is being said but also on who is saying it. When a talker becomes familiar (after sufficient listening time), listeners can build a stable model, thereby reducing acoustic uncertainty. Consistent with this view, speech perception is influenced by the listeners’ expectations about the talker, such as their dialect background (8, 9), ethnicity (10), accent (11), gender (12), and age (13). However, prior research has largely focused on coarse sociolinguistic contrasts, such as native versus non-native accents, leaving open whether subtler within-language variability, such as that encountered across multiple native speakers, induces acoustic uncertainty, which affects the generation and use of linguistic predictions.

Probabilistic expectations also play a central role in word recognition and sentence comprehension. As speech unfolds, lexical candidates that are compatible with the input encountered so far are incrementally pre-activated (4, 14, 15). For instance, upon hearing the phoneme sequence /kæp/, listeners may pre-activate lexical items such as *captain* or *capital*, with activation levels modulated by their relative likelihood in context, even before the full word is available. When additional information becomes available, lexical candidates are dynamically updated: some options are strengthened, while others are suppressed or ruled out entirely, allowing the listener to converge rapidly on the intended word. Moreover, listeners generate probabilistic predictions about upcoming words (3), drawing on context to predict not only word meanings (16–18), but also their grammatical structure (19–21) and even word form (22–25). Together, these findings support a hierarchical view of language comprehension in which graded predictions at multiple levels interact to support efficient processing (26). However, a key unresolved question is whether uncertainty at lower levels of processing, such as variability in the acoustic realization of phonemes, propagates upwards to constrain high level predictions. In the present study, we directly tested this possibility by examining to what extent talker variability modulates predictive processing during naturalistic speech comprehension.

Electroencephalography (EEG) signals were recorded from Italian native participants as they listened to continuous narrative speech under two conditions. In a Single-talker condition (hence Single) the story was narrated by a single talker, allowing listeners to build a stable, talker-specific model of the acoustic signal over time. This reduces acoustic uncertainty because the talker’s acoustic patterns become predictable. In a Multi-talker condition (hence Multi) the story was segmented and narrated by 9 different talkers, introducing substantial acoustic variability and reducing talker specific adaptation. Crucially, all talkers used canonical Italian pronunciation, allowing us to isolate the effect of acoustic variability independently of sociolinguistic differences. To link neural activity to speech and linguistic representations, we used the multivariate Temporal Response Function (TRF), a method that characterizes how specific stimulus features are encoded in time-resolved brain responses (3, 27–31). This approach provides an ecologically valid framework for examining how the human brain deals with acoustic variability during natural listening. Prior research using TRF has shown that speech processing involves the encoding of both low-level spectrotemporal information and higher-level categorical linguistic representations, such as phonetic articulatory features and phonemes (31, 32). Building on this work, our study examined how the brain copes with variability introduced by multiple (unfamiliar) speakers while simultaneously tracking multiple levels of representation. No previous study investigated how the brain tracks phonemic information as it unfolds over time despite differences in speaker-specific acoustic properties, alongside predictive mechanisms operating at both the lexical and sentential levels.

To model prediction during word recognition, we used phonemic surprisal which quantifies how unexpected a phoneme is given the preceding phonemes within a word (27). To capture prediction at the sentence or discourse level (i.e., between-word prediction), we used semantic surprisal which quantifies the degree of unexpectedness associated with a word given the preceding words (33). This measure is typically estimated via computational models such as large language models (e.g., GPT) (27, 37). The TRF-modelled neural response (weights) of phonemic surprisal typically exhibits fronto-central activity emerging within 350 ms after phoneme onset (27, 34), whereas the TRF-modelled neural response of semantic surprisal reveals a complex with prominent centro-parietal negativity, akin to the classic N400 component associated with prediction error (46, 53–55). Together these measures allow us to test how variability at the acoustic-phonetic level interacts with predictive processing across the linguistic hierarchy.

In our study, we employed linear modeling to investigate how talker variability affects the neural responses to phonemes (probing speech perception), as well as to phonemic surprisal and semantic surprisal (probing phonemic and semantic prediction, respectively). In the Multi condition, increased acoustic uncertainty derived from talker variability may demand greater processing of speech input to recognize phonemic categories, leading to enhanced neural responses to phonemes compared to the Single condition. Increased uncertainty in phoneme perception may also weaken the predictions about upcoming phonemes during word recognition, leading to reduced responses to phonemic surprisal. Finally, if acoustic uncertainty hinders the (pre-)activation of lexical-semantic representations, we would expect reduced responses to semantic surprisal in the Multi condition. Conversely, if uncertainty is resolved at the earlier processing stages, lexical-semantic predictions may remain unaffected.

## 2. Results

### 2.1. Phonemic processing

We used GAMMs (Generalized Additive Mixed Models) to assess condition differences in TRF responses to phonemes, averaging responses across phonemes and channels, as phoneme processing typically evokes temporally distributed neural responses (31). Model-based contrasts indicated significantly larger responses in the Multi than in the Single condition (smooth Time*Condition: *F* = 4.793, *p* < .001) within two time windows: an early positive peak [10, 200] ms, and a late negative deflection [270, 570] ms (Figure 2). These windows were defined as intervals where the 95% confidence interval of the smooth difference excluded zero (see Supplementary Material for complete model outputs and plot illustrating the same pattern for the phonetic features).

**Figure 1.**
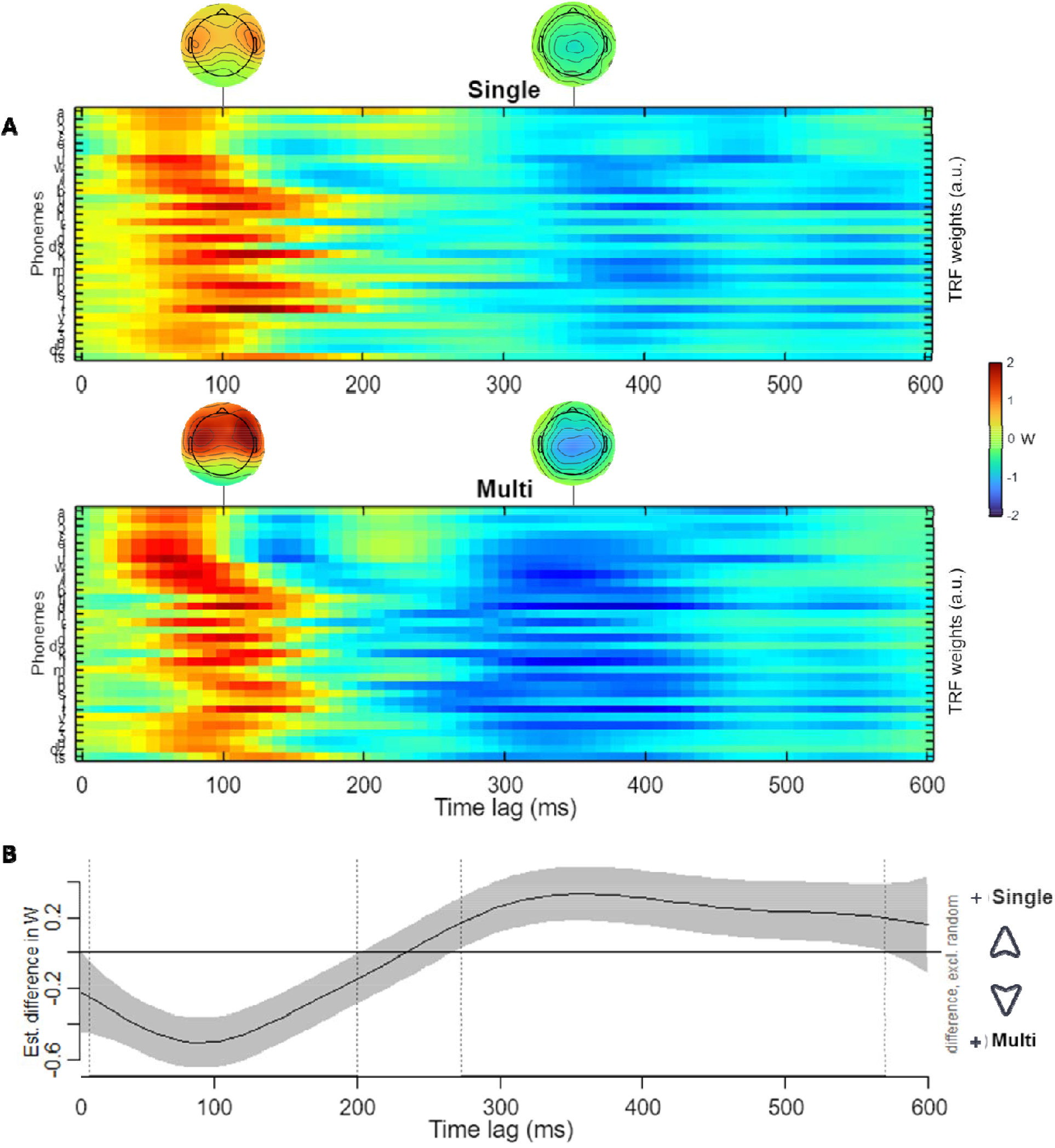
The brain promotes phonemic categorization to cope with acoustic uncertainty. Participants show a stronger response to phonemes in the Multi than in the Single condition. **(A)** Mean TRF weights of the phoneme regressor by Talker condition (Single above and Multi below) for all channels at post-stimulus time latencies from 0 to 600 ms. Each row represents the temporal fluctuation of TRF response to individual phonemes. **(B)** Estimated difference in TRF weights between the Multi and Single conditions (Single minus Multi) as a smooth function of time, obtained from a GAMM. Shaded areas indicate the 95% confidence interval. Time windows where the confidence interval does not include zero indicate reliable condition differences. Values above zero indicate stronger negative responses, whereas values below zero indicate stronger positive responses in the Multi compared to the Single condition.

**Figure 2.**
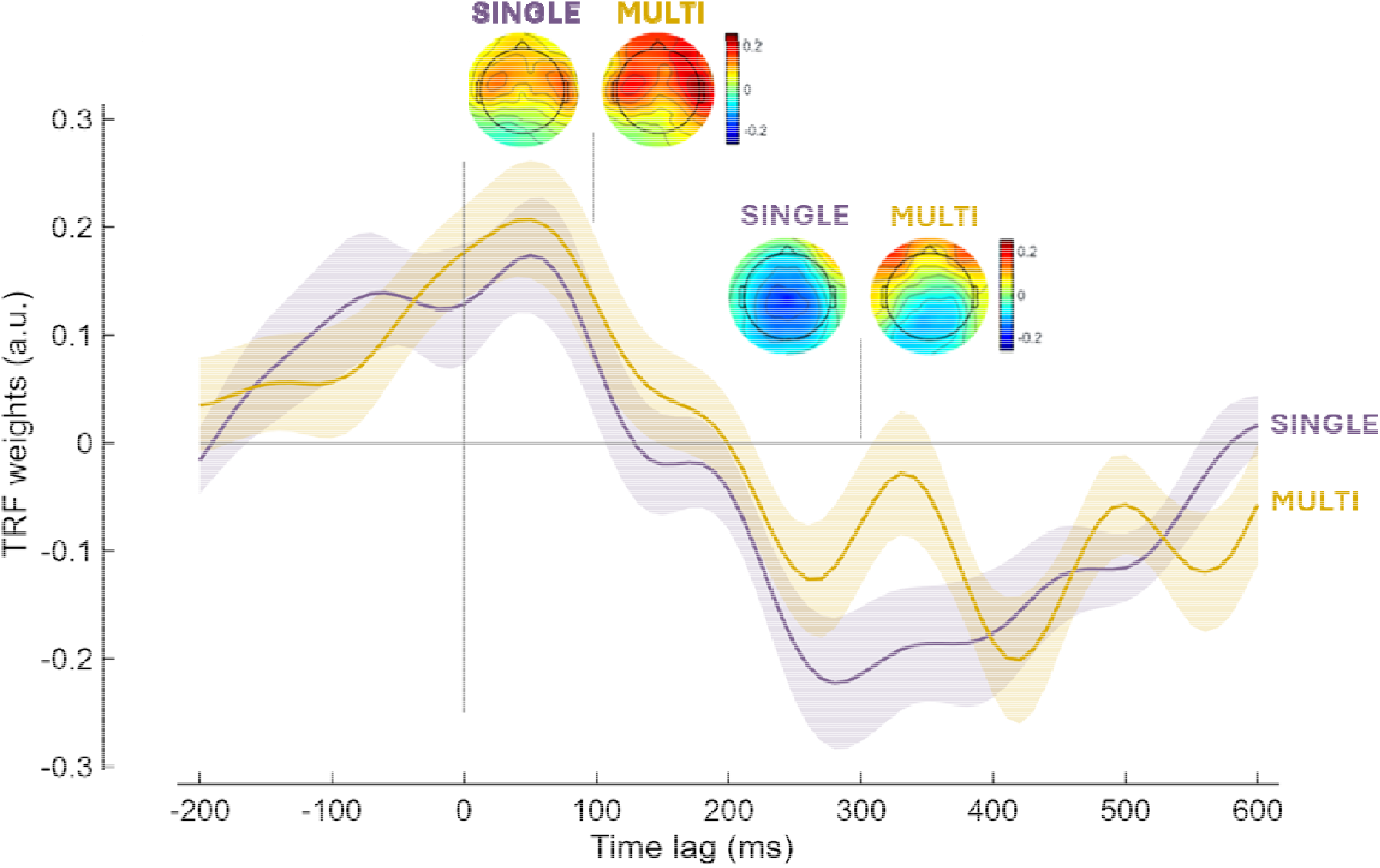
Acoustic uncertainty leads to reduced phonemic predictions. Average TRF weights of the phonemic surprisal regressor by Talker condition (Single and Multi) for Pz channel at post-stimulus time latencies from -200 to 600 ms. Shaded lines indicate SEM (Standard Error of the Mean) across participants. The figure also shows the scalp distribution of TRF weights for the phonemic surprisal regressor in the Single and Multi conditions at 100 ms and 300 ms.

### 2.2. Phonemic surprisal

We used Linear Mixed-effects models to test whether TRF responses to phonemic surprisal differed between conditions within the 0–350 ms time window, across a broad set of electrodes (F1, Fz, F2, FC1, FCz, FC2, C1, Cz, C2, CP1, CPz, CP2, Pz). Phonemic surprisal elicited a stronger negative response in the Single than the Multi condition (β = 2.125, *t* = -4.914, *p* < .001) (Fig. 2), indicating reduced phonemic prediction under increased talker variability. Note that the effect is significant also considering the [100, 350] ms window, thus excluding the first positive peak (β = 1.505, *t* = -4.588, *p* < .001).

### 2.3. Semantic Surprisal

We used Linear Mixed-effects models to assess whether TRF responses to semantic surprisal differed between conditions within the canonical TRF-N400 time window [300, 500] across a centro-parietal cluster of electrodes (FC1, FCz, FC2, C1, Cz, C2, CP1, CPz, CP2, P1,Pz, P2, POz). Semantic surprisal elicited a significantly larger centro-parietal negativity in the Multi than in the Single condition (β = 3.042, *t* = 3.060, *p* = .005), (Fig. 3), indicating enhanced semantic prediction under increased talker variability. Together, these results reveal a shift from phonemic to semantic predictive processing under increased talker variability.

**Figure 3.**
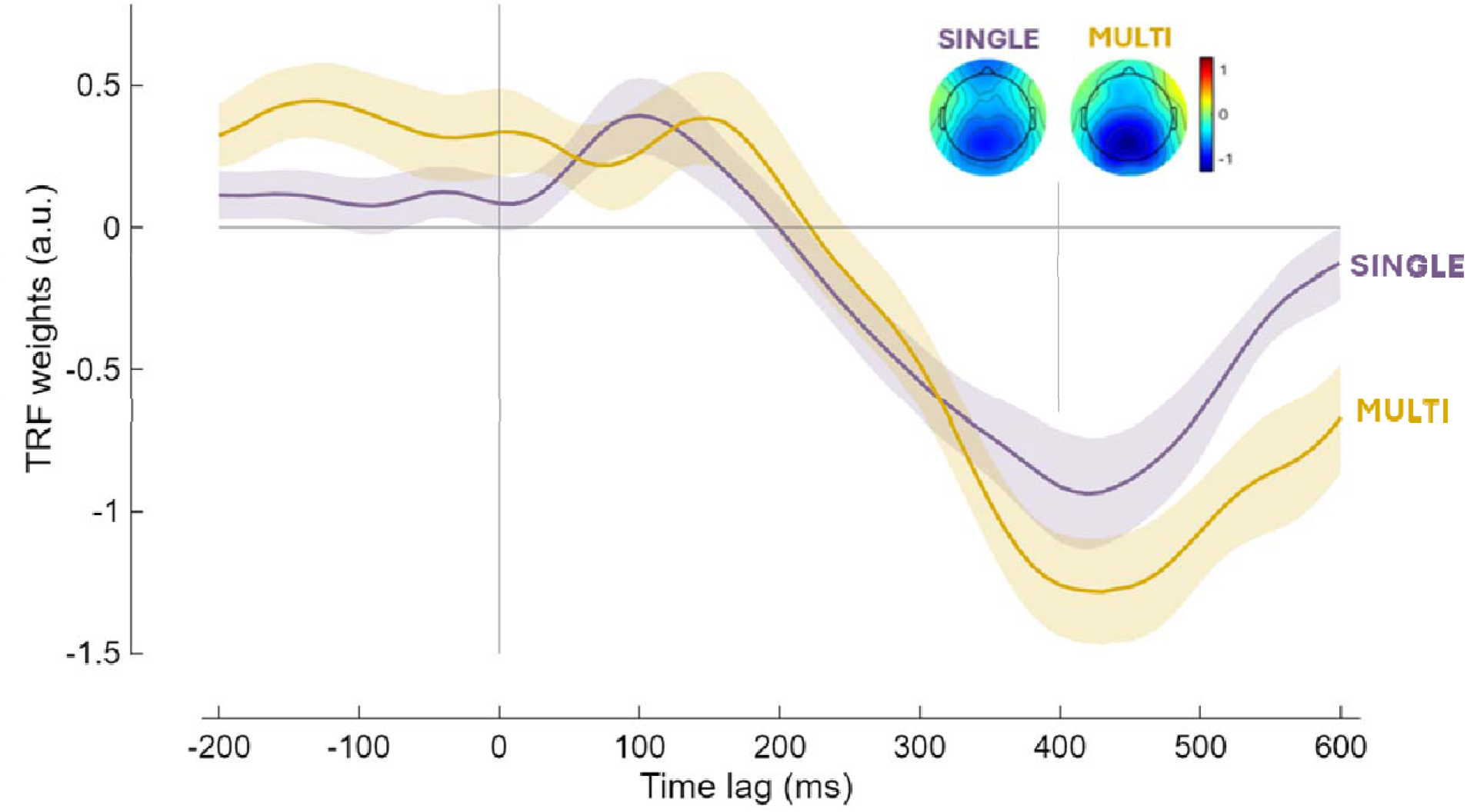
The brain relies on contextual semantic predictions to resolve acoustic uncertainty. Average TRF weights of the semantic surprisal regressor by Talker condition (Single and Multi) for Pz channel at post-stimulus time latencies from -200 to 600 ms. Shaded lines indicate SEM across participants (on Pz). The figure also shows the scalp distribution of TRF weights for the semantic surprisal regressor in the Single and Multi conditions at 400 ms.

## 3. Discussion

We show that natural talker variability reshapes predictive processing during speech comprehension, reducing phonemic prediction while enhancing semantic prediction. Increased talker variability prevents listeners from accumulating talker-specific experience, sustaining acoustic uncertainty that would otherwise be resolved through adaptation. This pattern reveals that predictive processes are not uniformly influenced by acoustic variability, but are flexibly redistributed across levels of representation. Rather than globally degrading prediction, increased acoustic uncertainty selectively weakens lower-level anticipatory processes while promoting reliance on higher level contextual information.

At the level of speech perception, we observed stronger neural response to phonemes in the Multi compared to the Single condition. This finding suggests that increased talker variability imposes greater computational demands on the mapping of speech sounds onto abstract phonological categories. When the entire story was narrated by a single talker, participants could accumulate talker-specific information and build a stable model of the acoustic-phonetic signal, facilitating speech perception. In contrast, exposure to multiple talkers gave listeners less time to adapt with each voice and therefore reduced the reliability of their internal model(s) (38, 39). The observed pattern aligns with adaptive accounts of speech perception, such as the Ideal Adapter Framework (5), which proposes that listeners continuously update probabilistic models of acoustic cues based on talker-specific experience. Crucially, our results extend this view by showing that even variability among native talkers modulates phonemic encoding during naturalistic listening, in line with previous results showing that language comprehension is shaped by how talkers adjust their speech across speech registers (37).

This interpretation is further supported by evidence linking adaptation to changes in low-level acoustic processing. The acoustic envelope provides an index of low-level sensory processing, reflecting sensitivity to acoustic properties. Recent evidence showed a reduction in neural encoding to acoustic envelope after the first 20 seconds, observed across multiple datasets, and time-locked to the onset of each experimental trial (reflected in both prediction correlations and P2 magnitudes; (40)). We argue this attenuation emerges from perceptual adaptation, after listeners have established a stable acoustic model, such that reduced neural encoding reflects more efficient processing of the acoustic input.

To directly examine how listeners use talker-specific information to facilitate speech perception, we conducted additional analyses comparing neural encoding of the speech envelope during the first and last 28 seconds of each trial, yielding approximately four minutes of data per participant (sufficient for envelope TRF estimate (41)). Results revealed within-trial adaptation in the Single condition, manifested as reduced encoding strength over the course of each trial. No such reduction was found in the Multi condition (an opposite trend was observed). This dissociation suggests that adaptation to a specific talker, rather than continuous listening per se, drives more efficient acoustic processing (see Supplementary Material for detailed results).

The present findings also provide compelling evidence that the effects of talker variability extend beyond speech perception, as reflected in systematic differences in phonemic and semantic prediction across conditions. At the level of phoneme sequences, increased acoustic uncertainty reduces neural responses to phonemic surprisal, indicating weaker top□down predictions about upcoming phonemes during word recognition. This reduced reliability increases bottom□up sensory demands, reflected by enhanced phoneme responses, while attenuating predictive coding, reflected by diminished surprisal responses. Because the linguistic content was identical across conditions, this effect cannot be attributed to differences in surprisal itself, but instead reflects how acoustic uncertainty influences the use of predictive information. This interpretation aligns with evidence showing that listeners wait for additional information from the incoming sensory input in the case of phonological errors produced by a foreign-accented talker (25).

In contrast semantic-level processing showed the opposite pattern. We observed enhanced neural responses to semantic surprisal in the Multi condition, suggesting increased reliance on contextual predictions about upcoming words. This finding indicates that when lower-levels processing becomes less reliable, listeners compensate by strengthening higher-level predictions. Such a shift is consistent with accounts of language processing in which multiple representational levels operate in parallel and can differentially contribute to comprehension depending on the reliability of the input (42, 43). In this context, semantic information may provide a more stable source of constraint, allowing listeners to maintain efficient comprehension despite increased acoustic variability.

Together, these results reveal a trade-off in predictive processing across levels of representation. Increased acoustic uncertainty reduces the strength of phonemic predictions while enhancing reliance on semantic expectations. This pattern suggests that predictive processing is dynamically reweighted rather than globally disrupted: when bottom-up information becomes less reliable, the system reallocates resources toward higher-level sources of information that can better constrain interpretation. Importantly, this trade-off indicates both interaction and partial independence between levels of the predictive hierarchy. While higher-level predictions can compensate for uncertainty at lower levels, the reduction in phonemic prediction shows that lower-level processes are not fully overridden by top-down expectations. These findings have broader implications for theories of predictive processing in language. Much of the existing evidence for hierarchical prediction has been obtained under relatively stable acoustic conditions, where variability is minimized. Our results demonstrate that under more naturalistic conditions, predictive mechanisms are sensitive to the statistical structure of the input and adapt accordingly. This supports a view of language comprehension as a flexible, context-dependent process in which the balance between bottom-up and top-down information is continuously adjusted.

In sum, we provide evidence that the brain dynamically reallocates predictive resources across levels of representation in response to acoustic variability. When talker variability increases uncertainty in the speech signal, phonemic prediction is reduced, but semantic prediction is enhanced. This flexible redistribution allows listeners to maintain robust comprehension in complex and variable listening environments, highlighting the adaptive nature of predictive processing in natural speech.

## 4. Methods

### 4.1. Participants

A total of 34 native talkers of Italian, aged between 18-35, were recruited to take part in the experiment (Female = 21). Of the original sample, 4 participants were excluded due to technical problems leaving the final cohort to 30 (Female = 19; M_age_ = 24.1 y.o., SD = 3.52). Participants were recruited from volunteers and students at the University of Padova, provided their informed consent before participating in the experiment, and were paid 25 euros for their participation. The research adhered to the principles outlined in the Declaration of Helsinki. The research protocol was approved by the Ethics Committee for Psychological Research of the University of Padova.

### 4.2. Material

Continuous Italian speech audios were employed in this study (see Data and Code Availability for stimuli and data). Two travel stories, each approximately 22 minutes long, were pre-recorded by male native Italian talkers (∼45 minutes in total; see Supporting Information for the transcription of the stories). The stories described trips to Ethiopia and North Korea, with content that was similarly novel to Italian participants. Each story was recorded twice: one for the Single-talker condition and one for the Multi-talker condition (Data and Code Availability for the audio recording files). In the Single-talker condition, the entire story was narrated by one talker, while in the Multi-talker condition, each section of the story was narrated by a different talker (9 talkers in total). In the Multi condition, the same set of talkers was used across both stories, whereas in the Single condition, different talkers narrated the two stories to avoid making our results specific to any individual talker. Talkers were instructed to speak naturally, which led to variations in speech rate (thus duration) across sections. Since previous literature showed that speech rate affects neural encoding of speech (37, 44), we matched the duration of corresponding sections between the two conditions (e.g., Section 2 of the Ethiopia story in Single and Multi conditions). This was achieved by calculating the mean duration of each section pair and applying minimal time-stretching or shortening (∼5%) to align the durations, while preserving pitch. The words and number of syllables were identical across conditions, ensuring that duration adjustments roughly corresponded to speech rate normalization. Both stories were normalized to a consistent average level of 68 dB (see Data and Code Availability to check the audio sounds).

We ran a pilot study on Prolific (www.prolific.com) to collect subjective ratings of audio sounds qualities that potentially affect speech processing (without being the focus of our manipulation). A total of 212 native Italian talkers rated one 2-minute story section on four dimensions: Clarity, Expressiveness, Pleasantness, and Naturalness, using a 7-point scale (1 = Not at all, 7 = Completely/Extremely).

We employed Cumulative Link Mixed Models (CLMM) to compare the two experimental conditions across the evaluated dimensions (Clarity, Expressiveness, Naturalness, and Pleasantness). CLMM are statistical models designed for analyzing ordinal data, where the variables have ordered categories and the distances between the categories are not known (45). These models estimate the probability that a response falls into a particular category or below, taking the ordinal nature of the data into account. Results showed no significant differences between conditions for any of the rated dimensions (all *p*-values > .05; see Supporting Information for visualizing the empirical distribution of the ratings on the evaluated dimensions).

Lastly, other acoustic features may differ across talkers in a naturalistic communicative scenario (e.g., pitch, phonemic realization). We deliberately did not control for such acoustic features, which naturally varied across talkers and were part of our manipulation to investigate adaptation to various talkers’ acoustics. To assess differences in the acoustic salience of speech sounds over time, we modeled non-linear fluctuations in the speech envelope (0-500 ms after each word onset) using Generalized Additive Mixed Models (GAMMs; 53). The model did not reveal a significant difference in amplitude between the two conditions across the analyzed time window (see Supplementary Materials). No analysis on the cohort entropy or semantic surprisal levels was performed as the two conditions shared identical words and phonemes (counterbalanced across stories and participants).

### 5.3. Procedure

All EEG data were collected in dimly lit and sound-proof booths. Stimuli were presented at a sampling rate of 44,100 Hz, monophonically, and at a comfortable volume throw loudspeakers. Participants were asked to listen attentively to two stories while EEG signals were recorded. They were asked to sit upright while looking at a fixation cross, which was presented on the centre of a computer screen right in front of them (at ∼80 cm of distance from their eyes). During the experimental session, participants were presented with one story per condition, with counterbalanced order across participants. To avoid any effects derived from specific relations between stories and conditions (e.g., a certain story is more interesting), each story was presented in both Single and Multi conditions across participants. The continuous narration of each story was divided into nine consecutive shorter trial of ∼2’30’’ minutes each. At the end of each trial, participants were asked 5 true/false comprehension questions (45 questions per story). Experimental sessions lasted ∼2 hours including preparation and testing.

### 5.4 Equipment

Electroencephalography (EEG) data were recorded using a 64 Ag-AgCl electrodes standard setting following the 10-20 international system (two actiCAP 64-channel systems, Brain Products GmbH, Germany) with hardware amplification (BrainAmp DC, Brain Products GmbH, Germany). Signals were bandpass filtered between 0.05 and 500 Hz, digitised using a sampling rate of 1000 Hz, and online referenced to the left mastoid. PsychoPy 2024 Software (47) was employed to present the stimuli. In addition, to ensure perfect synchronisation between the EEG recordings and the audio sounds, we implemented the Focusrite Scarlett 4i4 sound card (3rd gen) and the TriggerBox hardware (BrainProducts) to send triggers (see Supporting Information).

### 5.5. Analyses

#### 5.5.1. Behavioural data

Behavioural data were analyzed to identify and discard those participants with very low accuracy, who did not pay a sustained level of attention throughout the experiment. Each question could be scored either 0 to 1, which respectively represented wrong and correct answers. All participants scored above chance (∼80% on average, ranging between 67% and 93%). We did not further investigate behavioral data.

#### 5.5.2. EEG pre-processing

EEG signal analyses were performed on MATLAB Software (MathWorks, 2024b), using custom scripts, Fieldtrip toolbox functions (48), EEGLAB (49), and CNSP resources (50). Offline, the data were re-referenced to the average of the two mastoids, resampled to 100 Hz, and band-pass filtered between 0.5 and 8 Hz with a Butterworth zero-phase filter (order 2+2). Noisy channels with variance 3 times larger or smaller than the channels median variance were recalculated by spline interpolating the surrounding clean channels in EEGLAB. We had planned to exclude participants from the analysis if they had more than 30% of rejected data or more than six contaminated electrodes in over 30% of sections. However, no participants or sections were discarded for these reasons. Out of 18 sections, one story section for ten participants and two sections for two participants were excluded due to recording failures or contaminated data.

#### 5.5.3. EEG analysis

The cortical encoding of speech in the Single vs Multi conditions was estimated by measuring forward models or Temporal Response Functions, capturing the linear relationship between continuous stimulus features and the corresponding neural response. TRFs were calculated with the mTRF-Toolbox (29), which implements a linear regression mapping multiple stimulus features to one EEG channel at a time. The regression included an ridge regularization with parameter λ, and was solved through the closed formula *β*=(*X*^⊤^*X*+λ*I*)^−1^*X*^⊤^*y*, where β indicates the regression weights, *X* the stimulus features, *I* the identity matrix, and *y* an EEG channel. The interaction between stimulus and recorded brain responses is not instantaneous, as a sound stimulus at time *t*_*0*_ can affect the brain signals for a certain time-window [*t*_*1*_, *t*_*1*_*+t*_win_], with *t*_*1*_ ≥0 and *t*_*win*_ >0. The TRF takes this into account by including multiple time-lags between stimulus and neural signal, providing model weights that can be interpreted in both space (scalp topographies) and time-lag (speech-EEG latencies). Larger TRF weights (or linear regression weights), whether positive or negative, indicate that a particular feature at that time point plays a significant role in predicting the EEG signal. The time window used to fit the TRF model was [-200, 600] ms, based on previous research that found this time-lag window to be sufficient to capture the measurable EEG response associated with most cognitive processes underlying language comprehension (35, 37). To account for differences in scale and statistical structure across feature types, we used banded ridge regression, allowing separate regularization parameters for different feature groups. Specifically, distinct λ values were optimized for non-sparse regressors (e.g., acoustic envelope and spectrogram) and sparse regressors (e.g., onset and surprisal predictors). The regularization parameter was selected through an exhaustive search on a logarithmic parameter space from 10^−6^ to 10^6^. This selection was carried out via cross-validation to maximize the EEG prediction correlation averaged across all channels, leading to TRF models that optimally generalize to unseen data.

Leave-one-out cross-validation procedure was employed to maximize the amount of data used for the model fit, at the cost of additional computational time compared with a single train-test split. Stimulus and EEG time-series were split into nine folds, each corresponding to one of the nine segments of the stories. For each candidate λ value, the model was trained on all folds except one, which was used for testing. This process was repeated until each fold has served once as the test set. Each iteration provided a prediction correlation coefficient (*r-*value) between each feature and the EEG response (per channel). The prediction correlation coefficient is the estimate of how strongly an EEG signal encodes a given set of stimulus features. An *r*-value of 1 would represent perfect correspondence between EEG signal and TRF features, whereas an r-value of 0 would indicate no correlation whatsoever.

#### 5.5.4. Encoding of speech features (TRF regressors)

Given that complex stimuli such as speech typically include several dimensions, the multivariate form of TRF (mTRF) enables the simultaneous modeling of neural responses to multiple stimulus features. Here, we considered different predictors to investigate different levels of speech processing. We fitted a first model including acoustic spectrogram and phonetic features as predictors, which was used to probe phoneme perception. A second model included speech envelope, phoneme onset and phonemic surprisal, and was used to assess the extent to which listeners generate predictions about upcoming phonemes. The third model included speech envelope, word onset, and semantic surprisal, and was used to examine predictive processing at the lexical-semantic level, specifically with respect to upcoming words. By doing this, we aimed to investigate neural responses to different linguistic features, while accounting for acoustic differences across talker conditions (32, 37, 51). EEG and (non-zero) stimulus-vectors were normalized prior to fitting the TRF model (29, 50).

##### Acoustic envelope and spectrogram

The broadband acoustic envelope was extracted from the speech sounds using the Hilbert transform (30), which captures a key acoustic property of the speech material (52). The spectrogram regressor was implemented in 8 frequency bands ranging between 250 Hz and 8000 Hz. Those bands were defined based on the Greenwood equation that correlates the position of the hair cells in the inner ear to the frequencies that stimulate their corresponding auditory neurons (53). The spectrogram provides a richer and more detailed representation of the acoustic signal, capturing frequency variations essential for distinguishing phonemes and specific phonetic features (31, 54). Consequently, spectrograms are better suited to control for low-level acoustic information when probing phoneme perception.

##### Phonetic features (and phonemes)

Phonetic alignments of the speech material were obtained through forced alignment, initially performed automatically using WebMAUS (55) and subsequently verified manually with PRAAT (56). The alignments were stored as 17-dimensional binary time-series, where the different dimensions corresponded to phonetic articulatory features. Phonetic features indicated whether each phoneme was voiced, voiceless (consonants), plosive, fricative, affricate, nasal, approximant, front, back, central, close, open (vowels), bilabial, labiodental, dento-alveolar, palatal, velar-glottal. This way, each phoneme could be described as a particular linear combination of phonetic features. A linear transformation matrix was derived to describe the linear mapping from phonetic features to phonemes, which we used to rotate TRF weights from phonetic features to the phoneme domain (31, 32, 37).

##### Phonemic surprisal

Phonemic surprisal quantifies how unexpected a certain phoneme is given the sequence of phonemes listened so far. This measure is defined as the inverse of the conditional probability of each phoneme within a word, commonly approximated using corpus frequency data (27). In our study, word frequencies were extracted from ItWac Corpus (57), while their phonetic transcriptions were extracted from the WikiPronunciationDict, a multilingual pronunciation dictionary that includes Italian. Phonemic surprisal was calculated using a tree-based approach (27, 58). By constructing a probabilistic tree structure from the corpus, we were able to model the likelihood of various phoneme sequences and compute surprisal values at each phoneme position within words. We used WebMAUS (55) to force-align the phonemic transcriptions with the actual phoneme onsets in the audio stories, which we then manually verified. Surprisal values were subsequently assigned to the onsets of the corresponding phonemes.

##### Semantic surprisal

This metric quantifies how unexpected a word is in a given context and is computed as the negative logarithm of the lexical probabilities of the current word. For investigating encoding of semantic surprisal, we extracted word probabilities from the Italian transformer-based large language model UmBERTo (59). It is a model based on BERT and trained on large Italian Corpora (cased model trained on Commoncrawl ITA (∼69GB) exploiting 110M parameters and 32k vocabulary size). UmBERTo calculated the probability of each word of the story chuck given its context. Surprisal was given as the negative logarithm of the lexical probabilities of the current word. Surprisal values were then coded into a sparse time-vector, where non-zero entries represent word onsets, and their values the surprise of that word based on the context.

### 5.6. Statistical analysis

#### 5.6.1. TRF model performances

Before testing whether our manipulation influenced the TRF weights in the features of interest, we conducted a series of control tests to verify that these features were encoded in the neural signal in both conditions. Using a leave-one-out procedure (54, 60), we computed prediction-correlation gain for each participant and regressor (averaged across channels) and tested against zero using one-sample, one-tailed t-tests.

For phonetic features (modeled relative to the spectrogram), gain was significantly greater than 0 in both the Single condition (*t* = 6.104, *p* < .001) and the Multi condition (*t* = 6.336, *p* < .001). For the semantic regressors, prediction correlations were computed within reduced time windows corresponding to the expected latency of the neural response ([300 500] ms), as prediction-correlation metrics computed over the full model time-lag are known to dilute temporally localized effects (as described by 49). Both word onset (Single: *t* = 4.457, *p* < .001; Multi: *t* = 2.982, *p* = .006) and semantic surprisal (Single: *t* = 3.414, *p* = .001; Multi: *t* = 3.088, *p* = .004) regressor were encoded in the brain signals.

##### Isolating prediction-related activity

Δ*TRF-weights analysis*. Conversely, neural responses to phonemic information and onsets have strong temporal overlap (58, 61), which makes unique contribution of phonemic onset and surprisal difficult to disentangle within a windowed prediction-correlation framework (49). To address this limitation, we first assessed that phonemic surprisal and onsets together were significantly encoded in the brain signal (Single: *t* = 5.894, *p* < .001; Multi: *t* = 9.582, p < .001). Then, we adopted the ΔTRF-weights procedure proposed by (49), which directly contrasts TRF weights associated with surprisal and onset regressors. Then, surprisal-specific encoding was quantified by subtracting onset-related TRF weights from surprisal-related TRF weights and comparing this contrast against a null distribution obtained by shuffling the surprisal regressor non-zero values. This approach preserves the overall temporal structure of the data while isolating prediction-related activity beyond shared onset-driven responses. ΔTRF-weights were evaluated within a [0 350] ms time window, corresponding to the temporal range in which onset-driven and surprisal-driven responses are expected to differ (27). In both conditions, the difference between surprisal and onset response was significantly greater than the shuffle-based responses (Single: β = 0.216, *t* = 3.188, *p* = .002; Multi : β = 0.361, *t* = 5.584, *p* < .001), confirming a unique contribution of phonemic surprisal to the EEG signal beyond stimulus onsets.

#### 5.6.3. TRF regression weights

To investigate condition differences in feature-specific neural encoding, we analyzed the TRF weights of each regressor of interest. For semantic and phonemic surprisal regressors, we conducted window-based analyses focusing on time intervals motivated by prior literature. Phonemic surprisal effects were assessed within a [0–350] window, whereas semantic surprisal effects were evaluated within the [300–500] ms time window, consistent with previous TRF literature on phonemic and semantic predictive processing (3, 28, 35, 54, 60).

For each regressor and condition, we computed the area under the curve of the TRF weights within the relevant time window averaged across a priori defined electrodes of interest. Based on previous studies (3, 60, 62), the electrode selection reflected the largely overlapping yet subtly distinct topographies associated with semantic and phonemic processing: semantic surprisal was assessed over a central-parietal cluster (FC1, FCz, FC2, C1, Cz, C2, CP1, CPz, CP2, P1, Pz, P2, POz), whereas phonemic surprisal was assessed over a slightly more frontal cluster (F1, Fz, F2, FC1, FCz, FC2, C1, Cz, C2, CP1, CPz, CP2, Pz). This yielded one summary value per participant which was submitted to linear mixed-effects models (LME (63)) with Condition as a fixed effect and Participant as a random intercept. This approach allowed us to test condition-related differences in response amplitude while accounting for between-subject variability.

The temporal characteristics of TRF responses to phonemes are less consistently defined in the literature. Phoneme processing often elicits distributed temporal responses (31), with effects emerging between early auditory components (∼100 ms) and later negative deflections (∼300 ms). Consequently, differences between conditions are more likely to be reflected in fluctuations of the response over time, rather than in isolated peak amplitudes. A data-driven approach was therefore adopted to capture these dynamic response patterns. We characterized such temporal dynamics of the two conditions in the average TRF response across phonemes and channels ([0 600] ms) using generalized additive mixed models (GAMMs (64)). GAMMs model nonlinear relationships by estimating smooth functions, making them well suited for identifying temporally extended or non-monotonic differences in TRF responses. In these models, time was included as a smooth term, with separate smooth trajectories estimated for each condition, whereas condition was included as a parametric fixed factor, allowing for intercept differences. To account for individual variability, we included subject-specific smooth deviations from the group-level trajectory as well as random intercepts for subjects within each condition.

## Supplementary Material

Materials, data and analysis scripts will be made publicly available on OSF upon acceptance of the manuscript.

## Acknowledgments

G.P. was supported by a postdoctoral fellowship within the PRIN grant “Perceiving and predicting multisensory speech: a window on the interplay between sensory and motor processes and brain representations” (project code 2022FT8HNC), funded by the Italian Ministry of University and Research and awarded to F.P. This project has also received funding from the European Union’s Horizon 2020 research and innovation programme under the Marie Skłodowska-Curie grant agreement No. 101204985, awarded to G.P. M.S. was supported by a PhD grant from the University of Padova (2022–2025).

## CRediT author contributions

GP: Conceptualization, Investigation, Data curation, Formal analysis, Visualization, Writing - Original Draft; MS: Conceptualization, Investigation, Data curation, Visualization, Writing - Original Draft. RG: Investigation, Writing - Review & Editing. MMW: Resources, Methodology; FP: Conceptualization, Supervision, Funding acquisition, Writing - Review & Editing.

## Conflict of interest

The authors declare no competing interests.

## Notes

### Competing Interest Statement

The authors have declared no competing interest.

### Summary of Updates

We updated the analyses in the manuscript and report the revised results.

