## Supplementary Materials for "Adaptive Prediction: The Brain Trades Phonemic for Semantic Expectations Under Acoustic Uncertainty"

**This PDF file includes:**

Supporting text (T1 to T6)
Figures S1 to S3

**T1. Supporting Information Text**

**EEG-Audio Synchronization Setup Description.** The audio stimuli consisted of stereo sounds, where the left channel delivered the auditory stimulus to the participants, and the right channel contained a square wave marking the onset of each stimulus. Using a Focusrite Scarlett 4i4 sound card, we routed the left channel output to the loudspeakers and the right channel output to a TriggerBox device. Specifically, the right channel's square wave signal was connected to a BrainProducts StimTrack, which detected the onset pulses and forwarded trigger signals to the TriggerBox (see Figure S1). The TriggerBox then converted these signals into numeric triggers sent to the recording computer marking the start of each trial and ensuring precise synchronization with EEG recordings.

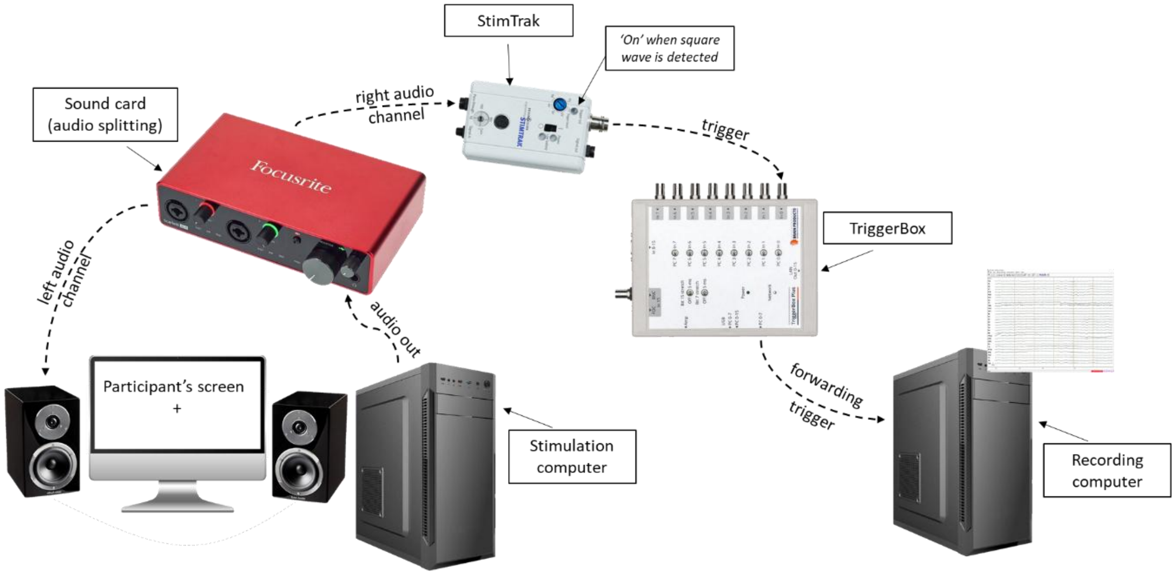


**Fig. S1.** Setup schematic representation.

**T2. Material's Subjective Rating Description.** Before the start of the experiment, we ran an online pilot study to collect subjective ratings of audio sounds qualities that potentially affect speech processing (without being the focus of our manipulation). A total of 212 native Italian speakers rated one 2-minute story section on four dimensions: clarity, expressiveness, pleasantness, and naturalness, using a 7-point scale (1 = Not at all, 7 = Completely/Extremely). Figure S2 shows the empirical distribution of the ratings on the evaluated dimensions.

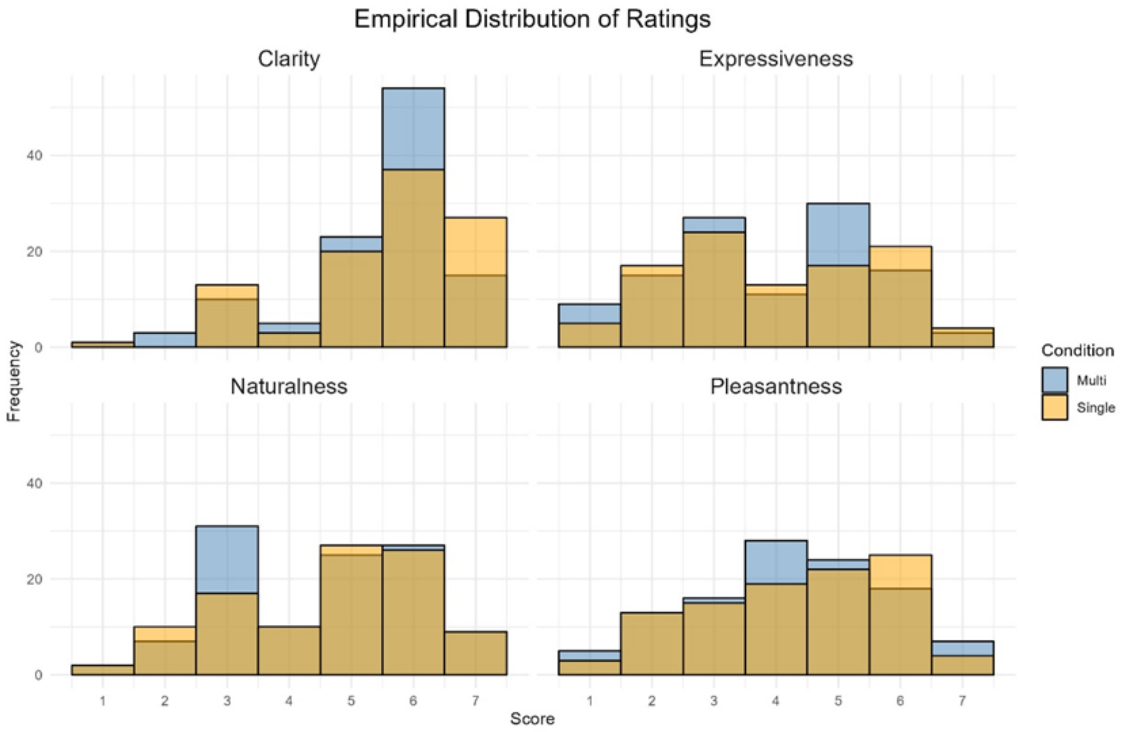


**Fig. S2.** Empirical distribution of participants' ratings of the narratives a 7-point Likert scale (1 = Not at all, 7 = Completely/Extremely) across four dimensions: Clarity, Expressiveness, Pleasantness, and Naturalness. Ratings are shown separately for the two experimental conditions, "Multi" (blue) and "Single" (orange). Each panel displays the frequency of scores within each dimension.

**T3. Stimuli Speech Envelope Analysis**
To assess differences in the acoustic salience of speech sounds over time, we modeled non-linear fluctuations in the speech envelope (0-500 ms after each word onset) using Generalized Additive Mixed Models (GAMMs). This approach allows the modeling of non-linear relationship between continuous variables (e.g., speech envelope amplitude and time). Specifically, we fitted a GAMM using the “bam” function of “mgcv” package with envelope amplitude as the dependent variable. The model included: (i) a main effect of Condition (Single vs. Multi) to test average amplitude differences between conditions, (ii) a smooth term for Time to estimate non-linear fluctuations of the speech envelope over time, (iii) a Condition-specific smooth over Time to assess whether non-linear envelope fluctuations differed across conditions, and (iv) time-smooth interactions for Speaker, Section, and Story to account for random non-linear variation across these grouping variables. The main effect of Condition was not statistically significant ($b = 0.01$, $t = 0.53$, $p = .60$), indicating no evidence of an overall difference in envelope amplitude between conditions. A significant smooth effect of Time ($F = 38.52$, $p < .001$) indicated non-linear fluctuations in the speech envelope within the 0–500 ms time window. However, the Condition-specific smooth over Time was not statistically significant ($F = 0.04$, $p = .85$), providing no evidence that the conditions differed in the envelope non-linear fluctuations over time.

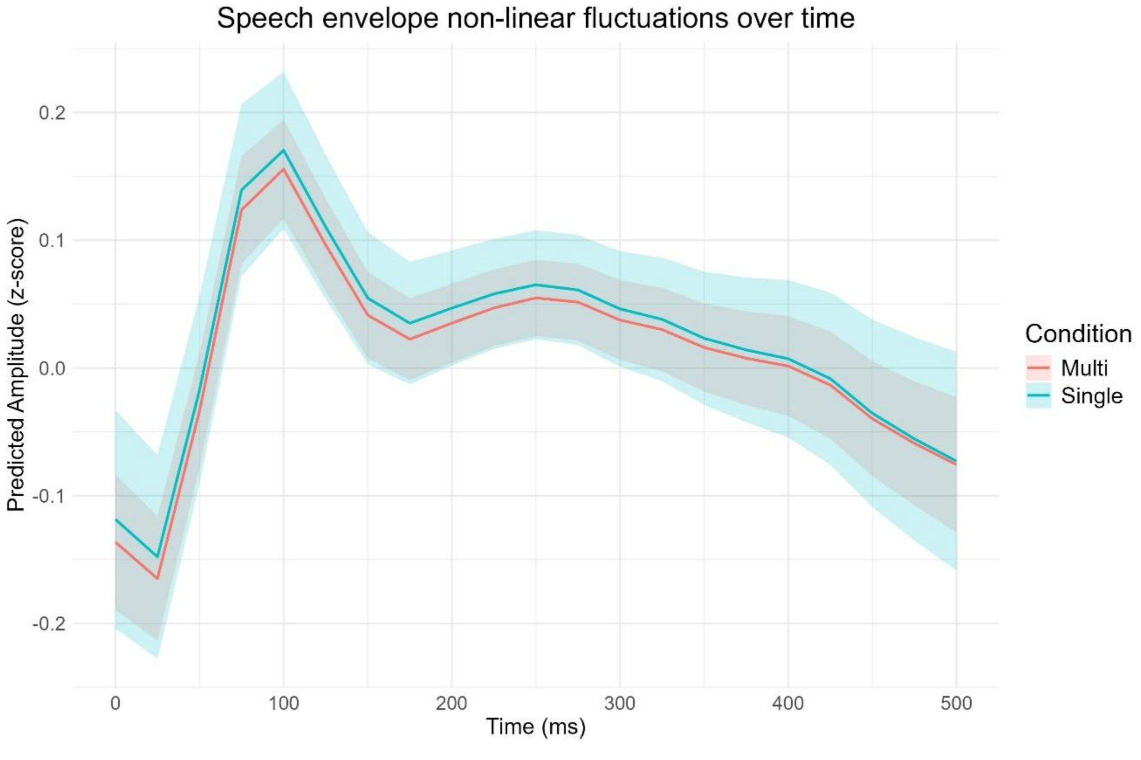


**Fig. S3.** GAMM Predicted fluctuation of stimuli speech envelope by condition (expressed as z-scores).

**T4. Speech Envelope TRF: Before-After**

Our results are consistent with the view that perceptual adaptation is a continuous, dynamic process: listeners progressively leverage talker-specific information to refine internal representations of speech sounds over time.
To further test this, we analysed our data by comparing the first and last 28 seconds of each block, yielding approximately 4 minutes of data per participant. Considering 30 participants, this quantity of data is probably sufficient to train a reliable envelope TRF model but remains likely underpowered for estimating sparse regressors (e.g., phonemic surprisal or categories). We expected to observe a reduction in envelope tracking within blocks of the Single condition, reflecting activation of the talker-specific model. In contrast, in the Multi condition, we expected the opposite pattern, as listeners must continuously adapt and update talker-specific acoustic models. As expected, we found that in the Single condition, speech envelope TRFs derived from the last 28 seconds of each trial showed reduced P2 magnitude ($\beta$ = 0.155, $t$ = 2.789, $p$ = .007) and lower prediction correlations ($\beta$ = 0.134, $t$ = 2.141, $p$ = .042) compared to the first 28 seconds. In contrast, these effects did not reach significance in the Multi condition, although the pattern was numerically reversed with slightly increased P2 amplitudes and prediction correlations in the last 28 seconds of each block (i.e., after more exposure to the same talker: see Supplementary Material for more details). These results may indicate not only low-level acoustic adaptation but also a facilitation in mapping the acoustic signal onto linguistic categories.

**T5. Additional Regressor (TRF Model).** In addition to the regressors included in the multivariate TRF model reported in the paper, based on UmBERTo, we computed semantic dissimilarity by measuring the cosine distance between the word embedding and the embedding of its preceding context. We added this regressor in the initial analysis to account for semantic integration on top of lexical prediction. However, this regressor's individual contribution was not significantly > 0 and was therefore excluded from further analysis.

**T6. Original Stories (Italian)**

**North Korea.**

1. Il mio viaggio nella Corea del Nord inizia con verdi viali di salici che delimitano strade a otto corsie dove transitano pochissime automobili, marciapiedi lindi e aiuole fiorite, quotidianamente curate da gruppi di cittadini volontari. È uno scenario unico fatto di architettura monumentale e celebrativa non imbrattata da cartelloni pubblicitari. Coreografiche manifestazioni di piazza ogni giorno in tutto il paese. Fiumi dalle acque limpide e pescose, pianure di campi di riso e mais, montagne tondeggianti ricoperte di boschi e interrotte da torrenti fruscianti. L'aria è pervasa da melodie delicate, Nessun cellulare che trilla irritante mentre cenate. In questo paese l'istruzione è gratuita, così come la sanità. Il sostentamento alimentare di base è garantito. I servizi essenziali come gas, luce e acqua hanno costi irrisori. La delinquenza è pressoché inesistente. Gli studenti del doposcuola imparano gratuitamente musica, pittura, ricamo e così via, mentre per gli adulti ci sono corsi di lingue e conferenze su ogni disciplina tecnico-scientifica. Le attività sportive fanno parte della vita quotidiana quanto il lavoro e i grandi centri urbani dispongono di ogni genere di impianto sportivo. Si tratta di un sogno, un'utopia? No: benvenuti nella Repubblica del Popolo Democratico di Corea. Sin dall'ingresso nel Paese dei Koryo, la Corea (dal nome della dinastia che governò sulla penisola coreana e che la unificò), ci si rende conto che non ci si trova in un posto qualunque, uno dei tanti "non luoghi" turistici, indistinguibili tra loro per il conformarsi allo standard del mercato. L'aeroporto internazionale di Pyongyang ti accoglie semivuoto, con pochi aerei fermi sulla pista. La breve attività quotidiana si concentra esclusivamente in coincidenza con il volo da e per Pechino. Le formalità sono rapide e i controlli brevi. La pattuglia degli ospiti stranieri è attesa, di loro si sa già tutto: per entrare nel paese è necessario richiedere per tempo un invito ufficiale, ottenibile tramite pochissime agenzie autorizzate previo acquisto di un pacchetto turistico. Il margine di manovra è limitatissimo, riducendosi in pratica alla durata del soggiorno (limitata a una dozzina di giorni al massimo) e alla scelta se visitare in gruppo o per conto proprio gli stessi posti, negli stessi orari. Ed è bene anche leggere tutte le avvertenze e i consigli per il viaggio, così non ci si sorprende quando in dogana vengono sequestrati i cellulari (che saranno poi restituiti all'uscita dal paese). Si ha la sensazione di star per entrare in un altro mondo, un mondo per l'appunto unico e complesso, certo non familiare: quando il mondo era diviso in due blocchi, chi visitava l'uno o l'altro poteva forse provare certe sensazioni.

2. L'idea che deve accompagnare il turista è quella del rispetto assoluto dei costumi e dei modelli culturali di questo paese: per questo, come prima cosa, sarà seguito sempre da almeno due guide (così l'una controlla l'altra in modo da evitare eventuali variazioni di programma) e non gli sarà permesso uscire da solo dall' albergo. L'hotel più frequentato dai turisti stranieri nella capitale è di forte impatto visivo: una torre di 47 piani in un'isola in mezzo al fiume. Il tempo per annoiarsi o aver voglia di fare due passi comunque non ci sarà, il contingentato programma turistico comincia sin dalla prima sera, con un affascinante spettacolo. L'evento si svolge dal 10 agosto al 10 ottobre per tre sere la settimana allo stadio della capitale, ed è un'elaborata coreografia composta da decine di migliaia di figuranti che si alternano tra danze, parate militari ed esercizi funambolici. Un riassunto vibrante e tridimensionale della storia del Paese (la traduzione del nome dell'evento significa per l'appunto "O mio caro", riferito in questo caso all' amor patrio). Parte dal secolo scorso, ovvero dall'occupazione giapponese, e arriva ai giorni nostri, con particolare riferimento agli ultimi 65 anni, cioè dalla nascita del Partito del Lavoratori con l'ascesa di Kim Il Sung, il Grande Generale. Lo spettatore apprende che è stato il Grande Generale a sconfiggere i giapponesi nel 1945, e che da quella data la Repubblica del Popolo Democratico di Corea è rinata, con lo sviluppo prima dell'agricoltura e poi dell'industria pesante. Grande rilievo è dato all'idea di riunificazione con la Corea del Sud e all'amicizia con la Repubblica Popolare Cinese, il più importante alleato politico ed economico. Nei giorni successivi all'arrivo le guide avranno modo di spiegare con dettaglio il ruolo degli Stati Uniti nel determinare l'attuale assetto della penisola coreana. In particolare, si concentrano sulla visita alla nave spia americana, catturata nel 1968 e ora trasformata in museo galleggiante al centro di Pyongyang, esibita con orgoglio e riadattata a mezzo di propaganda. I nordcoreani si ritengono vittime dell'aggressione imperialista, tuttora in atto, spiegata ai turisti e alle scolaresche in visita al museo con ricostruzioni tridimensionali in scala delle battaglie del periodo della Guerra di Corea.

3. Il culto della personalità e la dinastia dei Kim è dimostrata dalle statue giganti in bronzo brunito o da murales che ritraggono il Grande Generale in ogni angolo del Paese. Talvolta viene rappresentato in compagnia del figlio e attuale capo dell'esercito, Kim Il Jong, ufficialmente chiamato anche Caro Comandante, circondato da bambini o nell'atto di arringare la folla o in visita a una fabbrica. La presenza paternalistica del comandante in mezzo al popolo è sottolineata dalla sua proclamazione a "presidente eterno", caso unico al mondo di mantenimento post-mortem della più importante carica istituzionale. Anche il turista deve rendergli omaggio indossando il vestito buono: in fila, insieme a una folla di coreani abbigliati a festa, siamo stati condotti nel suo mausoleo per inchinarci profondamente, dai quattro punti cardinali, alla sua salma mummificata. Non viene esclusa dal circuito della deferenza la madre di Kim Il Jong e moglie del Grande Generale: l'omaggio è questa volta un mazzo di fiori da deporre sulla tomba nel cimitero dei martiri della rivoluzione. Per sottolineare quanto anche il resto del mondo abbia compreso la grandezza dei due Kim, il programma prevede una visita alla raccolta dei 250 mila preziosi regali offerti al Grande Generale e al Caro Comandante nel corso dei decenni dai Capi di Stato e di governo di ogni angolo del mondo. Ce ne sono alcuni provenienti anche dall' Italia. I doni, tanto preziosi quanto kitsch, spaziano da intere collezioni di manufatti d'avorio dei Capi di Stato africani a elaborati oggetti intarsiati di provenienza orientale. Il tutto è custodito in due grandi palazzi-cassaforte relegati tra le montagne del nord, ornati di marmi di Carrara e con le maniglie in simil-oro tempestati di finti rubini. La nostra curiosità di sapere se c'era già un nuovo leader in pectore destinato a succedere al Caro Comandante non ha trovato risposta. Qualche giorno dopo il nostro rientro in Italia, sono state date due notizie in rapida successione: la presentazione in Cina del terzogenito del Caro Comandante e la sua improvvisa nomina a generale a quattro stelle, coronata dall'investitura ufficiale. Questi compare a fianco del padre alla grande parata militare in occasione del 65-esimo anniversario della fondazione della Repubblica. Per la prima volta siamo quindi di fronte a un esempio di socialismo dinastico, con la trasformazione della dittatura del proletariato in dittatura ereditaria. L'unica cosa non ancora certa è il soprannome del Caro Comandante: tra i più probabili "Brillante Compagno" e "Giovane Generale".

4. Un giorno, una delle nostre sobrie guide ci ha sorpreso chiedendoci di informarlo sulle notizie dal mondo trasmesse dalle televisioni straniere. Abbiamo così scoperto che la televisione coreana riporta uno stringato e – naturalmente – censurato riassunto delle notizie dall'estero solamente il sabato. Accendere la TV è sempre interessante, in tutti i Paesi del mondo, ma nella Repubblica del Popolo Democratico di Corea è illuminante: canzoni popolari che inneggiano al governo e all'esercito, telegiornali che raccontano la visita del Caro Generale a qualche fabbrica o l'inaugurazione di un nuovo murales, costituiscono gli unici argomenti del palinsesto. E la televisione non è neppure posseduta dalla maggioranza delle famiglie. D' altra parte, l'isolamento e lo stretto controllo delle informazioni sono il modo più efficace per controllare la società, il mezzo più semplice per preservare la purezza del pensiero dai pericolosi germi esterni. Le immagini che potrebbero rivelarsi una potenziale fonte di turbamento all'ordine costituito vengono preventivamente censurate: con mia grande sorpresa tra le foto che mi sono state cancellate (oltre a quelle dei mezzi militari) ce n' erano di innocue che ritraevano comuni operai al lavoro, quasi che le attività manuali non fossero adatte a rappresentare la società. L'accurato isolamento informativo è affiancato da una martellante propaganda ben visibile agli angoli delle strade, in particolare nelle città di confine, dove megafoni a tutto volume mantengono alta la tensione, alimentando i timori di un attacco straniero e chiamando all'unità contro eventuali attacchi nemici. Nella capitale, il Palazzo della Cultura, un enorme edificio a forma di pagoda contenente cinque milioni di volumi, mette in pratica il concetto nordcoreano di educazione culturale: qui si tengono ogni giorno numerosi corsi gratuiti di lingue estere, di materie tecniche e di storia. I volumi hanno in prevalenza carattere scientifico, mentre la letteratura straniera è piuttosto datata. Internet è ancora un tabù, e gli accessi alla rete sono pressoché inesistenti. I libri in lingua italiana presentati su mia richiesta consistevano in una rivista di agricoltura e nell'edizione italiana di un libro di matematica superiore. Siamo rimasti stupefatti nel constatare che la nostra guida, un professore di lingua e letteratura inglese, non solo non avesse mai letto, ma non conoscesse neppure l'esistenza della saga di Harry Potter.

5. I grandi viali della capitale sono pressoché privi di traffico, con l'esclusione di poche Mercedes nere con i vetri oscurati a uso ufficiale e degli ancor più rari minibus turistici. Nelle strade ci sono quasi esclusivamente biciclette (poche) e pedoni (tanti) allineati ordinatamente in lunghissime file alle fermate dei vetusti autobus che collegano il centro con la periferia. Di notte la città piomba nel buio: i lampioni ci sono, ma pochi vengono accesi a causa della cronica mancanza di energia elettrica. Infatti, vista dallo spazio la notte della Corea del Nord sembra un buco nero accanto al bagliore del ricco sud. La gente vive per lo più in enormi grattacieli di cemento, ingentiliti da colori vivaci, che lo stato concede a uso gratuito. La guerra contro il Giappone e, soprattutto, quella contro gli Stati Uniti, hanno raso al suolo le città, e sono pochi i quartieri che ancora conservano le basse case tradizionali. Nella capitale c'è una bella metropolitana, scavata a 60 metri nel sottosuolo e collegata a numerose altre gallerie che possono essere utilizzate come rifugio antiatomico. Le stazioni sono prive dell'indicazione del nome della fermata e sono decorate in stile liberty con mosaici che ritraggono scene della moderna capitale o di vita quotidiana. Si accede mediante scale mobili attrezzate con piccoli altoparlanti che diffondono musiche patriottiche. In città i negozi sembrano invisibili, privi di insegne o di vetrine affacciate sulla strada. Di gestione statale, sono tutti simili tra loro e vendono pochi generi di consumo senza alcun assortimento. Ben più frequentati sono i numerosi parchi municipali lindi e ben curati, affollati a ogni ora del giorno da gente che li utilizza per fare picnic o per suonare e ballare all'ombra degli alberi mangiando coni gelato. Nelle piazze, invece, ci sono ore stabilite per le esercitazioni dei cittadini in preparazione delle frequenti parate celebrative. Se invece passiamo a parlare del senso di comunità, della disciplina militare e dell'esercito nordcoreano, questo è numericamente il quinto al mondo, grazie anche a una leva della durata minima di tre anni. È un dato notevole se si pensa alla estensione geografica del Paese.

6. L'ideologia della supremazia dell'esercito, ideata dal Caro Generale, è il cuore della macchina del potere. L'esercito è del popolo e la sua presenza si vede ovunque, sia nell'esecuzione di lavori socialmente utili come la realizzazione di infrastrutture o di abitazioni sia nelle parate militari o nei compiti istituzionali. Questo doppio utilizzo di soldati raggiunge lo scopo di cementare il patriottismo e di allontanare dalla gente l'impressione di una forza oppressiva. L'idea dell'appartenenza alla comunità è stimolata e ricompensata anche nei bambini con una settimana di vacanza in bellissime e attrezzate colonie estive. L'idea è questa: A differenza dell'occidente in cui è premiata l'eccellenza del singolo mediante l'esaltazione delle qualità individuali a scapito dei meno abili, il sistema nordcoreano premia solo il gruppo. Se in una scolaresca almeno la metà dei ragazzi ottiene il massimo dei voti, allora andranno tutti in vacanza premio, altrimenti nessuno. I migliori devono dunque darsi da fare per aiutare i meno capaci se vogliono essere premiati. Verrebbe così disincentivato l'egoismo e la lotta del tutti contro tutti per il successo del singolo. Viaggiando in minibus per le autostrade, si notano degli alti pilastri in pietra o cemento a gruppi di cinque o dieci a ogni lato della carreggiata, che aumentano di numero e frequenza avvicinandosi alle spiagge o al confine con la Corea del Sud: lo scopo è di bloccare le strade in caso di invasione nemica, facendoli saltare. E non sono gli unici esempi visibili che mostrano come la Repubblica si senta minacciata da pericoli esterni: le belle spiagge del Paese sono tutte protette da reti elettrificate che tolgono non poca poesia ai riflessi verde smeraldo del mare e alle sabbie dorate. Ci ha messo del suo anche la Corea del Sud, la quale ha costruito un bel muro di otto metri d'altezza sul confine, ben visibile da Nord e nascosto invece da Sud con un terrapieno su cui cresce un innocente erbetta verde. Ma è vero che il Popolo Democratico di Corea è l'Impero del Male, come sosteneva l'iper-semplificazione ideologica di George Bush? In realtà, l'eterno gioco al predominio tra superpotenze, dove oggi quella emergente contrapposta agli USA non è più la Russia, ma la Cina, vede la Corea, sia quella del nord che quella del sud, lacerata dai contrapposti interessi strategici ed economici dei due imperi.

7. Pedine e ostaggio di un gioco più grande, le due Coree non sono affatto libere di perseguire l'unificazione, come vorrebbe il nord, e neppure la denuclearizzazione del territorio, come auspicano entrambe, almeno finché gli Stati Uniti non rinunceranno alle loro basi nucleari in Corea

del Sud, a poche centinaia di chilometri dalla Cina. Su questo fatto si possono nutrire pochi dubbi, almeno per il futuro prossimo, perché c'è in ballo il controllo militare di una delle aree a più alto sviluppo economico del pianeta. L'ultima frontiera che vediamo è quella con la Cina, con il treno che ci riconduce a Pechino. È un balzo nello spazio e nel tempo. La sponda destra cinese del fiume che divide le due nazioni ospita una città piena di grattacieli e centri commerciali. Contrasta in modo netto con la riva sinistra nordcoreana, dove le fioche luci non riescono a rischiarare neppure le facciate in rovina dei vetusti condomini e le fangose strade non pavimentate. I nostri passi fuori dal treno, sui marciapiedi della prima stazione cinese, sono incerti: ci sembra strano poter camminare da soli ed entrare addirittura in un bar a prendere un caffè. Il Paese di Mezzo, traslitterazione dell'ideogramma della Cina, è una società che si sta trasformando a un ritmo travolgente. I mutamenti si sono intensificati negli ultimi dieci anni, rivoluzionando l'industria, il commercio estero, il turismo e persino la moda. A partire da queste premesse, non è più possibile ignorare quello che accade in questa parte dell'Asia. Il boom economico del paese è stato reso possibile da una forte riforma in senso liberista iniziata negli anni Novanta che ha reso possibile gestire in proprio un'impresa e che ha iscritto il paese all'Organizzazione mondiale del commercio. Queste due vere e proprie rivoluzioni hanno segnato la fine di un'era, quella dell'economia pianificata, e sono state prontamente metabolizzate, rendendo evidente la volontà di superare un passato di chiusura per vedersi riconoscere un ruolo adeguato all'aumentata forza economica. La Cina è oggi la nona potenza commerciale e la terza potenza spaziale al mondo. L'altra faccia della medaglia è che si sta cancellando in modo indiscriminato un passato che si percepisce come non più adeguato a rappresentare il nuovo che avanza. Interi antichi quartieri di città storiche sono stati spianati dalle ruspe per edificare al loro posto uffici, banche e centri commerciali.

8. L'isolazionismo stesso della Cina, durato tre decenni, ha lasciato il posto a un'ansia di modernizzazione che tuttavia non riesce a far proprie analoghe innovazioni nel campo dei diritti.
Tornando al viaggio nella Repubblica del Popolo Democratico di Corea, questo offre un'esperienza unica e al contempo surreale. Durante la mia esperienza, mi hanno stupito le rigide regole imposte dal governo e il costante controllo esercitato sulla popolazione e sui visitatori stranieri. A mio avviso, da questo emerge un quadro complesso di un paese che si definisce socialista, ma che si basa su dinastia ereditaria e autoritarismo. L'osservatore occidentale può sentirsi attratto dalla curiosità di esplorare un territorio così chiuso e misterioso, ma al tempo stesso si trova di fronte a un'atmosfera soffocante, permeata dalla costante propaganda e dal culto della personalità dei comandanti supremi. Le strutture monumentali, le parate militari, le statue imponenti e la martellante retorica patriottica dipingono un quadro di grandezza e forza, ma dietro questa facciata si nasconde una realtà più complessa e problematica. La mancanza di libertà individuale, la restrizione delle informazioni e la costante minaccia di rappresaglie per chi osa mettere in discussione il regime sono elementi che gettano un'ombra sulla vita quotidiana dei cittadini nordcoreani. Tuttavia, questa esperienza offre anche l'opportunità di riflettere sulla complessità delle relazioni internazionali e sulle dinamiche geopolitiche che influenzano la regione. La situazione della penisola coreana, divisa tra nord e sud da decine di anni di tensioni e conflitti, rappresenta un nodo cruciale nel panorama geopolitico globale. Inoltre, l'occasione di confrontarsi con una realtà così distante e diversa dalla propria può portare a una maggiore consapevolezza e comprensione delle sfide e delle contraddizioni che caratterizzano il mondo contemporaneo. Il viaggio nella Corea del Nord può quindi essere visto non solo come un'opportunità per esplorare un territorio poco conosciuto, ma anche come un modo per interrogarsi sulle proprie convinzioni e percezioni culturali. In definitiva, il viaggio nella Repubblica del Popolo Democratico di Corea può essere un'esperienza illuminante e stimolante, ma richiede anche una mente aperta e critica per cogliere appieno le complessità e le contraddizioni di questo affascinante e controverso paese. Questa è stata una vacanza in cui ho imparato moltissimo. Non è facile viaggiare in Corea del nord, ma tutti i preparativi e le scomodità che si devono affrontare valgono assolutamente la pena.

9. Ci si potrebbe domandare come e quando è avvenuta la divisione della Corea in Nord e Sud. Questa separazione è stata il risultato di una serie di eventi storici che hanno avuto luogo alla fine della Seconda Guerra Mondiale e nel periodo successivo. Durante la conferenza di Jalta nel febbraio 1945, le potenze alleate decisero di dividere la Corea lungo il 38° parallelo, con l'Unione Sovietica che amministrava il Nord e gli Stati Uniti il Sud, in attesa di elezioni per un governo

unitario. Tuttavia, l'ideale di una Corea unita si dissolse rapidamente a causa dell'escalation delle tensioni tra le due potenze vincitrici, che rappresentavano ideologie e politiche contrastanti: il comunismo nel Nord e il capitalismo nel Sud. Nel 1948, la Corea del Nord proclamò la sua indipendenza come Repubblica Democratica Popolare di Corea, mentre la Corea del Sud fondò la Repubblica di Corea. La divisione fu ufficializzata dalla comunità internazionale, ma non rispettava le aspirazioni della popolazione coreana, che desiderava un'unificazione nazionale. Le tensioni tra i due regimi rivali portarono a scontri armati lungo il confine, culminando nella Guerra di Corea nel 1950, quando il Nord invase il Sud. La guerra durò tre anni e si concluse con un armistizio nel 1953, senza un vero trattato di pace. Da allora, le due Coree rimangono tecnicamente in stato di guerra, dal periodo della guerra fredda fino ad oggi. La divisione ha avuto conseguenze significative per entrambe le Coree. Il Nord ha abbracciato un sistema politico stalinista e isolazionista, con un'economia centralizzata e un culto della personalità intorno ai leader della dinastia Kim. Nel frattempo, il Sud ha seguito una via capitalista, diventando un'importante economia industriale e tecnologica con il supporto degli Stati Uniti. La divisione ha generato una serie di questioni irrisolte, inclusi i conflitti familiari, la separazione delle famiglie e la presenza di una zona demilitarizzata lungo il confine. Come è avvenuto per la divisione tra Germania Ovest e Germania Est, intere famiglie si sono trovate spaccate e divise contro la propria volontà. La zona demilitarizzata coreana è visibile di notte dallo spazio. È lunga 250 chilometri con circa 4 km di larghezza. Paradossalmente, sebbene la zona che separa entrambi i lati sia demilitarizzata, il confine oltre quella striscia è uno dei confini più pesantemente militarizzati del mondo. Nonostante gli sforzi per promuovere la riconciliazione e il dialogo inter-coreano, la divisione rimane una delle questioni geopolitiche più durature e complesse del mondo, con profonde implicazioni per la stabilità di tutta la regione circostante e la sicurezza internazionale.

**Ethiopia.**

1. Avevo timore di visitare la Dancalia in Etiopia, per via delle vecchie cronache di rapimenti e uccisioni di turisti nei pressi del confine con l'Eritrea. I viaggi turistici in quella zona sono ripresi solo nel duemila diciotto, dopo una sospensione durata un lustro. La storia del confine tra i due paesi che si sviluppa problematicamente nella prima metà del Novecento, è abbastanza complessa. Nel millenovecentosessanta due, l'allora regnante dell'Etiopia annetté l'Eritrea come provincia dell'impero etiope, con il tacito avallo delle potenze occidentali. Dopo trent'anni di resistenza armata, l'Eritrea ha ottenuto nel mille novecento novanta tre l'indipendenza dall'Etiopia in seguito allo schiacciante risultato di un referendum. Ma la faccenda non era destinata a risolversi tanto presto: da allora per lunghi anni, i due paesi sono rimasti invischiati in dispute sui confini. A causa dell'imprecisa definizione della frontiera stabilita nel mille novecento due fra l'Italia fascista, allora presenza coloniale in Eritrea, e l'Impero d'Etiopia, lo status della località e delle aree circostanti non fu mai del tutto chiaro. Per quella linea immaginaria morirono circa ottanta mila soldati, in uno scenario bellico devastante. Poi, dal duemila, con gli accordi di Algeri, si giunse a un cessate il fuoco provvisorio, ma i continui scontri tra le bande di ribelli eritrei e i militari etiopi hanno messo in pericolo l'incolumità dei turisti di passaggio. Così, nel duemila sette e poi ancora nel duemila dodici, si sono verificati episodi di violenza che hanno portato alla chiusura della regione al turismo per anni. All'inizio del duemila diciotto nessuno avrebbe immaginato che erano maturati i tempi per cambiamenti radicali nelle relazioni tra Etiopia ed Eritrea. Un fattore decisivo è stato l'ingresso sulla scena africana dell'ultimo decennio di un nuovo attore commerciale e politico, ovvero la Cina. La svolta politica è stata ufficializzata nel giugno di quell'anno, quando il presidente etiope ha dichiarato che il suo esecutivo avrebbe rinunciato alle rivendicazioni territoriali in Eritrea. Poco dopo, in luglio, è stata firmata una dichiarazione che ha posto fine allo stato di guerra tra i due paesi con la riapertura della rotta aerea diretta tra le due capitali, Addis Abeba e Asmara, del commercio bilaterale e delle rispettive ambasciate. In seguito alla firma degli accordi di pace è tornato relativamente sicuro visitare le zone di frontiera tra i due paesi, tra cui la depressione della Dancalia.

2. La depressione etiope a forma di triangolo nasce dalla divergenza di tre placche tettoniche nel Corno d'Africa, allontanamento che ha generato un bassopiano vulcanico e arido che si estende sotto il livello del mare. L'intero territorio è disseminato di conetti vulcanici ed è solcato da fenditure

e faglie tettoniche. L'etnia che popola al novanta cinque per cento la Dancalia oggi vive di pastorizia in piccoli villaggi dove l'acqua è fornita da autobotti del governo e parte del cibo da aiuti internazionali. La popolazione abita in capanne, tende o edifici precari. Si entra nel territorio previo pagamento di un biglietto e la visita richiede l'accompagnamento di una guida e di poliziotti di scorta. Presto, abbiamo iniziato a notare uomini armati lungo la strada, tra cui ragazzini che portavano con noncuranza un'arma a tracolla, oltre ai militari. È un retaggio del passato, che ora fa effetto, ma la militarizzazione delle strade qualche anno fa si giustificava per via del conflitto armato di cui sopra. Questa situazione, lungi dal trasmettere una sensazione di minaccia, non è mai stata fonte di tensione data l'assenza di atteggiamenti aggressivi unita a un'intenzione amichevole pur se distaccata. La maggior parte delle persone sono cortesi e sostanzialmente indifferenti mentre i bambini tendono a interagire più attivamente con i turisti. In aree come queste la scuola è logisticamente difficile da frequentare e i bambini passano molte ore a giocare per strada. L'impressione è che si muovano col tacito consenso degli adulti per cercare di carpire qualche moneta ai turisti di passaggio, improvvisandosi guide o vendendo fossili, l'importante è saper stare al gioco e dare mance adeguate. Siamo stati testimoni di ragazzini che hanno scagliato pietre contro i nostri fuoristrada, forse solo per gioco, forse per noia o passatempo. nonostante avessimo un poliziotto armato a bordo. La sua deterrenza sembrava valere poco o nulla. Comunque si è trattato di un episodio isolato. Se da un lato è triste pensare che questi ragazzini siano figli di una guerra recente, dall'altro lato si percepisce un po' di speranza per il futuro. La dura tradizione del passato è stata cancellata dalla fine della guerra e di un impero, oltre che dalla crescita demografica. Oggi le dinamiche sono altre, dietro alle filantropiche iniziative di sviluppo, finanziate e portate avanti dalla Cina nel sostanziale disinteresse dell'Europa, si nascondono forse nuove ambizioni neocoloniali.

3. Le nuove strade e ferrovie promuovono sviluppo e commercio, addomesticano e rimpiccioliscono il territorio. Come effetto collaterale finiranno presto col dare il colpo di grazia all'economia del sale trasportato a dorso di cammello, retaggio di un passato che ancora si rifiuta caparbiamente di scomparire. Impieghiamo un'ora per uscire da Addis Abeba attraverso una grande e moderna autostrada: arteria costruita con l'aiuto della Cina per i camion che vanno e vengono da Gibuti, ormai porto cinese di Addis Abeba. Adesso, grazie alla Cina e ai suoi capitali, la maggior parte delle vie non sono più sterrate e il manto stradale è di ottima fattura. Gli interessi del Dragone si toccano con mano anche nel pieno centro della capitale, dove è in costruzione un nuovo grande stadio di calcio. Appena ci si allontana dai luoghi di interesse, la strada presenta numerose buche e interruzioni dovute ai lavori in corso. La carreggiata spesso è invasa da animali vaganti che rendono pericoloso spostarsi dopo il tramonto, come ci spiega il nostro autista. Lungo questa parte di tragitto abbiamo contato diversi gravi incidenti con mezzi pesanti rovesciati che ostruivano la sede stradale. Dell'antico tracciato ferroviario restano solo alcune stazioni abbandonate e dai binari divelti. Ci siamo fermati al ristorante di una vecchia stazione, gestito da un affascinante signora di origine greca. L'anziana dal volto antico, inciso da rughe come una ragnatela, si è rivolta a noi in italiano e ci ha raccontato dei tempi andati, quando i treni passavano da qui, quando c'era la ferrovia. I tavoli del ristorante sono riparati da pergole che poggiano sulla pensilina deserta dove, chiudendo gli occhi e liberando la fantasia, non è difficile immaginare l'arrivo di nuvole di vapore, forti rumori metallici e il brusio dei lavoratori e delle persone in transito. È un luogo simbolico, dove il presente è sospeso tra la decadenza e gli ingombranti segni del passato coloniale. Se tutto andrà bene, ci faranno un parcheggio. Ma per un mondo che scompare, un altro avanza. La ferrovia tra Addis Abeba e Gibuti è stata ultimata nel 2016 dal Gruppo ferrovie della Cina e passa non lontano da qui, pur non toccando più il centro del villaggio. Per lunghi anni, nel corso della guerra, l'Etiopia non ha avuto uno sbocco sul mare.

4. Così, dopo i falliti tentativi di collaborazione economica dell'Unione Europea nei primi anni duemila, la Cina si offerse di finanziare e costruire una linea a scartamento normale, nell'ambito di un piano di costruzione di una vasta rete ferroviaria nell'Africa orientale. Oggi e ancor più in futuro i prodotti cinesi viaggeranno su rotaia. E la più diretta conseguenza sarà appunto l'estinzione del commercio del sale, scavato a mani nude dagli scalpellini nel cuore della depressione della Dancalia che molti millenni fa fu invasa dalle acque del Mar Rosso. Un mestiere durissimo, quello

del cavatore di sale, praticato a mani nude unicamente con l'aiuto di bastoni e scalpelli tra le prime luci dell'alba e le prime ore del mattino. Poi, quando il sole si alza verso lo zenit, le attività si fermano per riprendere prima del tramonto, quando le temperature tornano più sopportabili. E nonostante questo è possibile lavorare solo in certe stagioni. Prima che faccia scuro, i dorsi consumati dei cammelli sono caricati con le lastre squadrate. Venti lastre, all'incirca duecento chili per cammello, decine di cammelli per carovana. Ne abbiamo individuata almeno una composta esclusivamente di montoni. Sono le "carovane del sale", sale sporco, pieno di sedimento, grigio e marrone, destinato esclusivamente all'uso alimentare animale. I cammelli viaggiano perlopiù di notte per fermarsi a riposare all'ombra nelle ore più calde della giornata. Nel paesaggio etiope ci sono tanti elementi naturalistici di estremo interesse. Come, ad esempio, i due vulcani che sorgono rispettivamente a poca distanza da un lago e nel bel mezzo della depressione della Dancalia. Dal campeggio sul lago arriviamo in mezz'ora di auto al villaggio da cui partono le missioni per il primo vulcano. Vengo introdotto dopo lunga attesa presso un signore etiope di età per me indefinibile, né giovane né anziano, che giaceva placidamente sdraiato all'ombra, sulle stuoie della sua capanna, attorniato da quelli che a prima vista avevano tutta l'aria di essere dei luogotenenti. Lui, a quanto pare, è il solo a decidere chi deve accompagnarci nonché l'entità del contributo da versare. Tutti i gruppi che passano da queste parti, attirati dal vulcano come falene da una fiamma, devono passare di qui e confrontarsi con questa figura dall'aria autorevole ma forse dall'investitura non esattamente istituzionale. Per l'occasione in veste di coordinatore, sono io a sobbarcarmi l'onere e l'onore di essere ammesso alla sua presenza. Lo saluto cordialmente, con un certo grado di deferenza, e vengo fatto accomodare nella sua capanna, l'epicentro appunto di un potere non meglio definito. Sono ben conscio che da lui dipende il come e, soprattutto, il quanto della nostra salita ai vulcani.

5. La speranza è che un pizzico di cordialità possa portare ad un ammorbidimento tariffario. Speranza vana: la trattativa disastrosa si conclude con un alleggerimento della cassa di circa 15 euro a testa (in moneta locale) per il permesso di transito e di salita. Poco male, l'entusiasmo per aver staccato il biglietto immaginario per il vulcano è tanto e pone in secondo piano altre considerazioni di sorta, per il momento. In aggiunta, ci è assegnata una scorta armata di due poliziotti: ne basterebbe uno solo, ma ce ne fanno comunque pagare due. Per non farsi mancare nulla, guida obbligatoria, anche se i nostri autisti conoscono perfettamente la strada avendola percorsa innumerevoli volte. Faccio buon viso alla "trattativa" salutando il signore con un sorriso. C'è ben poco da trattare quando non si ha il coltello dalla parte del manico. Caricata la giovanissima guida imbocchiamo un terreno sterrato che serpeggia in mezzo a colate laviche recenti. Venticinque chilometri e due ore di scossoni dopo arriviamo al villaggio alla base del cammino, un agglomerato di capanne in pietra di grandi dimensioni dove solitamente i gruppi si fermano per attendere il buio. Nonostante l'ora e il caldo, decidiamo di salire subito per ammirare il tramonto dalla cima del vulcano e fare una prima circumnavigazione della caldera alla luce del giorno. Iniziamo la salita all'una e trenta del pomeriggio, dopo un pranzo leggero. Ce lo sconsigliano, ma il caldo secco sui trentacinque gradi centigradi risulta sopportabile, purché si cammini con lentezza e s'indossi un cappello largo. Chi ce l'ha usa un ombrellino come riparo aggiuntivo dai raggi del sole intenso. Percorriamo i nove chilometri e mezzo che separano il villaggio dal campo alto sul cratere in parte su strada e poi seguendo un largo sentiero. Impieghiamo tre ore esatte. Il dislivello in salita è di 400 metri. Anche se un cammello al seguito basta e avanza, le dimensioni del nostro gruppo impongono che ne prendiamo due, (due per due giorni) anche se le ore effettive della nostra permanenza sul vulcano risulteranno inferiori a ventiquattro. Infine, saldiamo l'uso al campo alto di tre capanne in pietra col tetto di paglia. La situazione abitativa non è comodissima, ma lo spettacolo attorno richiama allo spirito di adattamento.

6. Per farla breve, "l'operazione vulcano numero uno" richiede cinquanta euro circa a testa tra annessi e connessi. Arriviamo al campo alto in tempo per il tramonto e, con grande sorpresa, scopriamo di essere soli. Il pallido disco solare è inghiottito dalla foschia che aleggia bassa sull'orizzonte intorno alle sei del pomeriggio: abbiamo quindi tutto il tempo per fare il giro della caldera sommitale. La scelta si rivela felice perché riusciamo a vedere di più durante il giorno che nell'escursione notturna, quando il fumo e i vapori abbondanti fungono da ostacolo, senza contare che di notte ci ritroviamo in mezzo a decine di gruppi di turisti che vagano alla cieca sul bordo del

cratere come anime dannate nell'inferno dantesco. Il tramonto è magico, soprattutto perché abbiamo il vulcano tutto per noi. Come generoso compenso per la resilienza dimostrata durante le salite e gli accidentati tragitti, i turbinanti vapori che dall'aprile del 2018 precludono quasi del tutto la vista del lago di lava sottostante si aprono per un po'. Riusciamo a intravedere per un attimo il fiume di lava scorrere sul fondo, contrariamente a tutti gli altri che ci raggiungono in seguito. Dal primo vulcano impieghiamo un giorno intero per trasferirci nel cuore della depressione della Dancalia, dove si trova il secondo vulcano. Decisamente diverso dal primo questo è molto particolare: sembra di stare su un altro pianeta, un sito affascinante, che a mio avviso vale il viaggio in Dancalia. Una parte del fascino sta nelle sorgenti termali, risultato dell'esplosione di una camera magmatica nella valle posta sotto un importante deposito di sale lasciato dal Mar Rosso al suo ritiro. Oggi si presenta come una piatta distesa di incrostazioni multicolori: gialle, arancioni, verdi. Ossidi si accumulano nelle pozze, ognuna delle quali ribolle attraverso minuscoli geyser fumanti, circondati da un terreno che ha tutte le sfumature del marrone. Nell'aria aleggia un vago odore di petrolio. Passeggiando in questo paesaggio alieno, novanta metri sotto il livello del mare, che non riceve neppure duecento millimetri l'anno di pioggia e dove le temperature possono raggiungere i sessanta gradi, si ha l'impressione di trovarsi su Marte. Faccio attenzione a non calpestare i sottili merletti calcarei, composti da evaporiti di carbonati e sali di sodio e potassio. Chissà quanti secoli ci sono voluti per la loro formazione. Non ci sono transenne né recinti, chiunque può andare dove vuole.

7. Questa citata libertà comporta una responsabilità che a volte manca in noi turisti: temo arriverà presto il momento in cui spunteranno percorsi obbligati e recinti. Nonostante l'ostilità dell'ambiente, sono qui stati scoperti di recente, con il contributo di geologi italiani dell'università di Bologna, dei microrganismi che riescono a sopravvivere in ambienti che superano i cento gradi centigradi in condizioni super acide e sature di sale. Questi batteri dei geyser permettono di studiare quali siano le condizioni limite per la vita sulla Terra, estendendo il concetto di abitabilità anche ad altri pianeti. Adiacente ai geyser multicolori, c'è un luogo che vale la pena di visitare per lo stesso motivo per cui ci si reca nella vecchia stazione dismessa che citavo prima. È il villaggio fantasma degli italiani. Si notano qua e là, sullo sfondo di montagne azzurrine che si ergono oltre la pianura di sale, erosi dalla ruggine e quasi dissolti dal tempo, scheletri di camion Fiat, pezzi di binario, cisterne, una caldaia di locomotiva, una jeep, costruzioni diroccate di mattoni di sale. È l'insediamento della Compagnia Mineraria Coloniale, che qui ha scavato sin dal mille novecento 12. Controllata dalla Banca Italiana di Sconto, il cui principale azionista era Giovanni Agnelli, la compagnia finanziò la costruzione d'insediamenti minerari per l'estrazione del potassio. Il potassio, tra gli altri usi, serve anche nella preparazione degli esplosivi e per questo si può affermare che, chi riuscì a sfruttare la concessione durante la Prima guerra mondiale, ottenne un buon profitto. Nel periodo coloniale italiano i depositi furono sfruttati dalla compagnia mineraria dell'Africa orientale, il cui commercio partiva da un piccolo porto eritreo sul Mar Rosso. Con la fine dell'impero, gli inglesi eliminarono la ferrovia e smantellarono parte del villaggio italiano: senza l'accesso al mare, in una zona così difficile, l'estrazione mineraria risultava non proficua e il sito andò in decadenza. La Natura è ora libera di riprendersi i suoi spazi e le vestigie industriali stanno lentamente scomparendo, erose dalla ruggine e sepolte dalle sabbie come dinosauri, come se la Terra si volesse riappropriare di quel luogo inospitale e inadatto agli uomini, ma incantevole nella sua terribile bellezza. L'Etiopia è un luogo carico di ferite storiche e bellezza selvaggia. Attraversare questo territorio è un viaggio che porta il visitatore attraverso le cicatrici del passato coloniale, le sfide della modernizzazione e la resilienza della gente che vi abita.

8. L'apertura delle rotte turistiche nella regione segna una svolta importante nelle relazioni tra Etiopia ed Eritrea, offrendo l'opportunità di esplorare un territorio ricco di cultura e paesaggi unici. La presenza cinese, dopo quella europea, da un lato porta sviluppo e infrastrutture, dall'altro si confronta con tradizioni secolari e l'equilibrio ambientale della regione. Le comunità delle zone più remote vivono ancora in condizioni di estrema povertà, dipendenti dagli aiuti esterni. Le nuove infrastrutture portate dalla Cina potrebbero portare cambiamenti radicali nella vita di queste persone, sia positivi che negativi. La sfida per il governo etiope sarà quella di bilanciare lo sviluppo economico con la preservazione delle tradizioni e la tutela dell'ambiente. Il turismo potrebbe giocare un ruolo fondamentale in questo equilibrio, fornendo entrate economiche alle comunità

locali e promuovendo la conservazione culturale e ambientale. Le sfide future del governo etiope includeranno la stesura di leggi e normative per regolamentare le attività tradizionali della zona, come l'estrazione del sale, ma soprattutto il turismo ai vulcani. Ciò potrebbe implicare la creazione di norme per garantire la sostenibilità ambientale e la sicurezza delle persone coinvolte in tali attività, oltre a fornire formazione e supporto per migliorare le pratiche lavorative. Il destino delle persone che vivono vicino ai vulcani dipenderà in larga misura dalle politiche e dalle azioni del governo locale e nazionale. È importante che qualsiasi intervento tenga conto delle esigenze e dei diritti delle comunità locali, promuovendo uno sviluppo sostenibile che rispetti l'ambiente e migliorando al contempo le condizioni di vita delle persone coinvolte. In conclusione, l'Etiopia è un luogo di contrasti e meraviglie, dove il passato si intreccia con il presente e il futuro. È un luogo che merita di essere esplorato con rispetto e attenzione, per scoprire la sua storia, la sua gente e la sua bellezza unica. Spero di tornare presto a scoprire un altro pezzetto di Etiopia.

9. Nonostante recentemente nella regione sia tornata una situazione di relativa stabilità, le guerre passate e il colonialismo italiano hanno lasciato dei segni indelebili in Etiopia ed Eritrea. Molte persone sono fuggite da situazioni disastrose: alcune di loro sono emigrate in Europa e proprio in Italia. Questo ha portato ad un fermento culturale definito postcoloniale, che ha preso varie forme artistiche. Per esempio, si è espresso attraverso la letteratura, la musica oppure il cinema. Questo movimento ha portato, ad esempio, allo sviluppo della letteratura italiana 'postcoloniale', scritta e raccontata da emigrati in Italia, o da italiani di seconda generazione. Molti scrittori e scrittrici provenienti dal corno d'africa, tra cui le ex colonie italiane, quali Etiopia, Eritrea e Somalia, hanno vissuto sulla propria pelle la violenza del colonialismo italiano e delle guerre che si sono scatenate al termine del periodo imperialista. Le tematiche ricorrenti di questo filone culturale abbracciano le difficoltà e le tragedie legate all'attraversamento del mediterraneo da parte degli immigrati considerati clandestini. Un altro tema affrontato è la fine del controllo diretto delle potenze Europee che ha portato a una forte destabilizzazione nei governi della regione e a sanguinosi conflitti. Alcuni autori si focalizzano invece sulle politiche interne della regione. Un caso specifico è quello del colonialismo dell'Etiopia a discapito dell'Eritrea e della sua conseguente lotta per l'indipendenza. Questi casi mostrano che il colonialismo non è un fenomeno limitatamente europeo o italiano, ma potenzialmente di ogni forza prevaricatrice. Va sempre tenuto presente, però, che a monte di questi conflitti interni al corno d'africa (cosi come in altre regioni africane), il colonialismo e la ritirata delle potenze europee ha sicuramente avuto un peso nel creare forti squilibri tra nazioni africane. Le diverse forme di espressione di questo filone culturale raccontano esperienze di colonialismo nelle sue varie forme che guardano criticamente al mondo contemporaneo.
